# Exploring the impact of political regimes on biodiversity

**DOI:** 10.1101/2020.02.25.964429

**Authors:** Alexander Zizka, Oskar Rydén, Daniel Edler, Johannes Klein, Heléne Aronsson, Allison Perrigo, Daniele Silvestro, Sverker C. Jagers, Staffan I. Lindberg, Alexandre Antonelli

## Abstract

National governments are the main actors responsible for mapping and protecting their biodiversity, but countries differ in their capacity, willingness, and effectiveness to do so. We quantify the global biodiversity managed by different regime types and developed a tool to explore the links between level of democracy and other key socio-economic variables with the number of natural history specimens registered within country boundaries. Using this tool, distinct and previously unknown patterns emerge around the world, that urge for increased collaboration between the natural and social sciences to further explore these patterns and their underlying processes.

## Introduction

Political decisions often lead to differences in biodiversity research effort^1^ and the availability of biodiversity data.^2,3^ The mode of this decision making can vary from democratic to autocratic.^5^ A country’s level of democracy is determined by multiple dimensions, including suffrage (the fraction of citizens entitled to vote), quality of elections, freedom of expression and association, and constraints on executive power.^6,7^ Although these aspects co-vary to a certain extent, the significant variation across countries demonstrates that individual dimensions of democracy may affect the availability of biodiversity data (and also biodiversity conservation) differently. For this reason, it is misleading to use uni-dimensional indicators (e.g. simply democratic vs. autocratic) to explore the effect of political systems on biodiversity knowledge.^8^ More nuanced, numerically continuous measures of countries’ political institutions are key to understand why biodiversity observation data are collected and reported to varying degrees.

Varieties of Democracy (V-Dem) provides such data for the first time allowing us to more thoroughly explore how political regimes influence conservation efforts and the availability of biodiversity data via several mechanisms.^1^ For instance, compared to more autocratic regimes, liberal democracies generally have more open and reliable legal systems, which makes democracies more accessible for researchers and conservationists to collect and share biodiversity data.^9^ Free, fair, and regular elections provide citizens with a means to express their demand for good environmental conditions and thus also incentivize political leaders to invest in biodiversity management.^10^ In countries where the threat of conflict or physical violence is lower, fieldwork is safer, especially for international researchers.^11^ In addition, countries with higher levels of education may have a higher overall level of environmental awareness. This, together with a higher freedom of association, may lead to the development of ecological and naturalist societies,which contribute considerably to the availability of biodiversity data (including “citizen science”, which feed data into open platforms such as www.ebird.org and www.inaturalist.org).

We quantify the proportion of global biodiversity that is managed by different regime types^12^ and explore the relationship between the availability of primary biodiversity occurrence data (geo-referenced natural history specimens and species observations; obtained from www.gbif.org) and various democratic institutions, indexes and categories.^7^ Specifically, we ask three questions: 1) Which fraction of the global biodiversity is managed by democratic or autocratic regimes?; 2) How does the availability of primary biodiversity data relate to the political situation across countries?; and 3) What is the relation between democratization and the availability of primary diversity data through time? To address these questions, we calculate the range-weighted species richness of three vertebrate groups and seed plants per country globally, and develop a novel software (Bio-Dem, www.bio-dem.surge.sh), which is freely available.

## Results and Discussion

The analyses of distribution data from 22,805 vertebrate species and 333,986 seed plant species, show that the majority of the global species are managed under democratic regimes, mostly electoral democracies (Fig 1a). However, a few autocratic countries—–such as China, Venezuela, Madagascar and Papua New Guinea—report high biodiversity and hence are of critical importance for conservation (Fig. 1b). The electoral democracies in South America contribute to a disproportionately large share of global biodiversity under democratic rule (Fig. 1c). The recent autocratization and rise of populism in this region is concerning, potentially generating severe consequences for biodiversity conservation and the environment in general.^13,14^

**Figure 1:**
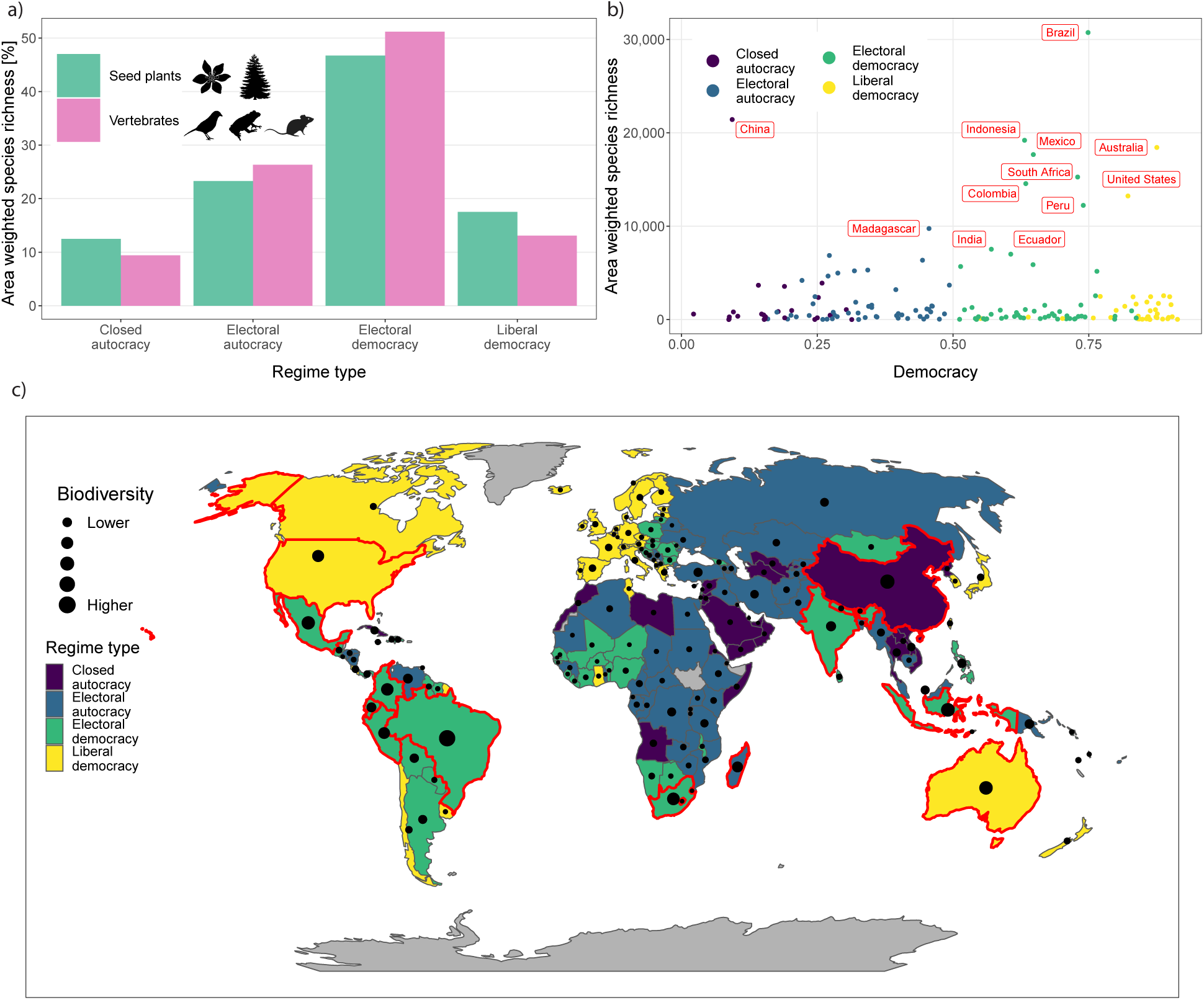
The majority of the world’s vertebrate and seed plant diversity is managed by democratic regimes. Biodiversity is approximated as range-weighted species richness of amphibians, mammals, and non-marine birds, democracy is measured by importance of elections in a country (polyarchy) and regime type. **a)** The majority of global biodiversity is managed by democratic countries, predominantly electoral democracies. **b)** The relationship between vertebrate diversity and level of democracy. **c)** The high fraction of biodiversity managed by democracies is mostly due to high biodiversity and levels of democracy in South America. Red labels and outlines in b) and c) point to the twelve most biodiverse countries globally.

Exploring the availability of biodiversity data in the context of political regimes around the world, reveals several distinct and hitherto poorly documented patterns. For instance, the amount of available biodiversity data correlates with electoral democracy (Fig. 2a). Similarly, the density of available biodiversity data mirrors the average number of years of education (Fig. 2b). Costa Rica emerges as an outlier, with a high density of occurrence records despite the country’s relatively low average education period. Conversely, numerous countries formerly part of the Soviet Union stand out by their low number of records but high average education length.

**Figure 2:**
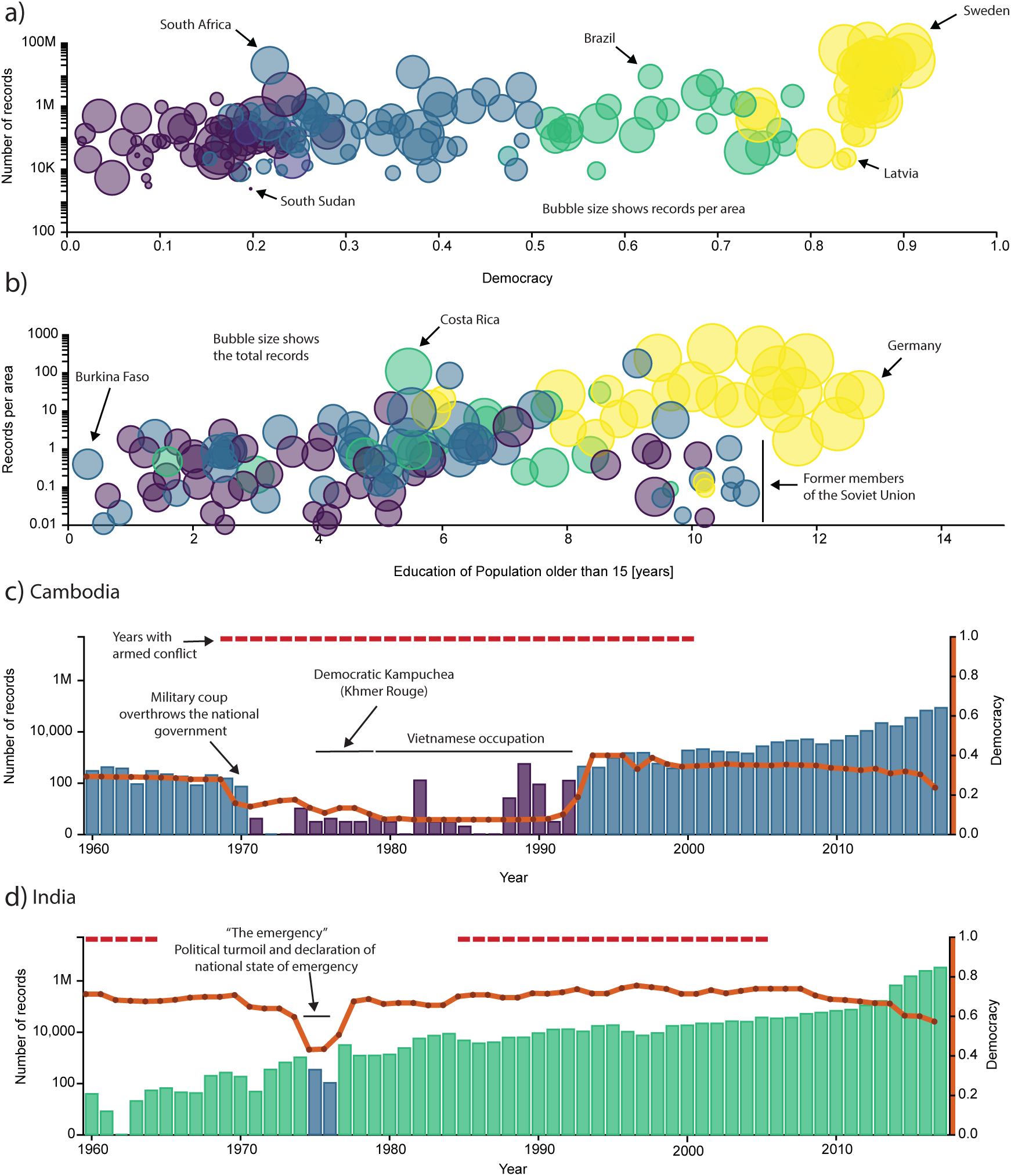
Biodiversity data availability reflects the state of political systems through time. Colours indicate regime type as in Fig. 1. **a)** There is no clear correlation between democracy and amount of area protected, but liberal democracies have generally more records available per area. **b)** Countries with a longer average period of education have on average more biodiversity data available. Bio-Dem allows the exploration of country-specific patterns: **c)** In Cambodia, a period of autocratization and armed conflict is related to a decrease in biodiversity data available between 1970 and 1992. **d)** In India, a period of political emergency and the resulting drop in democratic rights correlates with a drop in record availability from national institutions by one order of magnitude between 1975 and 1976.

Many countries change political regime type over the course of their history. Bio-Dem enables the exploration of how regime change, as well as armed conflict, affect the availability of primary biodiversity data. Taking Cambodia as an example, we find a decrease of new biodiversity data records by two orders of magnitude in the 1970s, coinciding with the beginning of a period of conflicts and autocratization (Fig. 2c). The end of this period and the corresponding increase in the level of democracy led to a sudden increase in data availability. Similarly, in India political turmoil and a related decrease in the level of democracy in 1975 and 1976 led to an abrupt decrease in the availability of biodiversity data from national institutions (Fig. 2d). Despite historical turmoil and a minor recent decline in the level of democracy, Cambodia and India mirror most other countries in exhibiting a general increase in biodiversity data, probably attributable to the widespread use of citizen science applications for mobile phones such as iNaturalist.

The relationships between political differences, socio-economic variables and biodiversity knowledge are multi-faceted.^1^ These links are also likely to be multi-directional. Increasing societal concerns for environmental protection are likely to affect political processes and the gathering of biodiversity data. In general, a causal interpretation of the observed patterns is difficult, due to indirect or unclear mechanisms, and the large number of potentially confounding factors.^1^ The Bio-Dem app and its underlying data sources provide a useful platform for research at a global and regional scale and over time. We hope it will foster increased collaborations between biologists, conservationists and political and other social scientists.^15^

## Material and Methods

Commented scripts for all analyses are available in the electronic supplement of this article and the source code of the Bio-Dem app is available at https://github.com/AntonelliLab/Bio-Dem under a MIT license.

We used two datasets of species geographic distributions to estimate the fraction of species covered by regime type. For amphibians, non-marine birds and mammals, we used publicly available geographic ranges from the International Union for the Conservation of Nature (www.iucn.org) together with country borders as provided by Naturalearth (www.naturalearth.org) to estimate the range-weighted species endemism per country.^16^ We first downloaded the ranges for all species, excluded marine birds based on expert knowledge (because most of their ranges are in international waters), and overlaid the range of each species with country borders. We then divided the size of a species’ range within each country by the total range size of the species and summed the values for all species per country. For instance, if a species is endemic to a country (i.e., the entire range is within country borders), it adds 1 unit to the country’s species richness, and if 10% of a species range is within a country this species adds 0.1. We then combined this per country species richness with data on species threat level (www.iucn.org) and with information on the state of democracy in each country for the year 2017 from V-Dem (https://v-dem.net)^7^ for the visualizations in Figure 1. For plants we approximated the species range with data from the World Checklist of Selected Plant Families (WCSP, https://wcsp.science.kew.org). WCSP provides distribution information on level-3 of the World Geographical Scheme for Recording Plant Distributions (www.tdwg.org). We used this information as the species range and continued the analyses in the same way as described for animals.

The results presented here were generated by novel software developed for this study, the Bio-Dem web application (www.bio-dem.surge.sh). Bio-Dem is a free app implemented in JavaScript. It allows users to explore the relationship between biodiversity data availability and the state of political regimes across countries globally and through time (since 1900). The app includes a large number of political as well as socio-economic indicators of expected relevance to biodiversity data collection and mobilization. It further allows faceting the data by time period and biological group. Bio-Dem obtains information on species occurrence records from the GBIF API and data on political indicators from the Varieties of Democracies database version 8 (www.v-dem.net). The app allows the generation of publication-level graphs in an easily accessible way. All data shown in Figure 2 are exported from Bio-Dem with minimal further editing.

## Acknowledgements

We thank Rafaël Govaerts for providing the plant data from WCSP. The research presented in this paper is a contribution to the strategic research area Biodiversity and Ecosystems in a Changing Climate, BECC. Funding was provided by iDiv/sDiv via the German Research Foundation (DFG FZT 118) to AZ; the Centre for Collective Action Research to OR; Riksbankens Jubileumsfond (Grant M13-0559:1), Knut and Alice Wallenberg Foundation (Grant 2013.0166) to SIL; internal grants from the Vice-Chancellor’s office, the Dean of the College of Social Sciences, and the Department of Political Science at the University of Gothenburg to SIL and SJ; the Swedish Research Council, the Knut and Alice Wallenberg Foundation, the Swedish Foundation for Strategic Research and the Royal Botanic Gardens, Kew to AA.

## Author contributions

AZ, AA, OR, SJ, DS, AP, and SL conceived of this study. AZ and HA analysed the data. AZ, OR, DE and JK developed the Bio-Dem application. AZ and AA wrote the manuscript with contributions from all authors.

## Notes

https://bio-dem.surge.sh

